# Targeted Memory Reactivation Increases Memory Recall: A Meta-analysis

**DOI:** 10.1101/796458

**Authors:** Thomas Lieber

## Abstract

Research on Targeted Memory Reactivation (TMR) shows that it’s a successful for increasing memory recall. However, As of yet no systematic study had been conducted to evaluate to what extend TMR can increase recall. This study used a comprehensive literature search to attempt to find all studies conducted in which TMR is used to increase memory recall. Methodological quality, study characteristics and where possible descriptive statistics were extracted from the studies. In total, 41 studies were identified, 26 of which included enough descriptive data to be included in the meta-analysis. The results indicated that the average methodological quality was good. Furthermore, TMR significantly increased memory recall. The subgroup analysis for modality was not significant, while the subgroup analysis for type of memory did yield a significant result. Overall, TMR seems to have small to medium effect on memory recall, is independent of modality and depends on memory.

## Introduction

Targeted memory reactivation (TMR) is a method whereby cues associated with previous learning are used to externally reactivate aspects of this learning (Oudiette & Paller, 2013). TMR is based on the idea of hippocampal replay during slow wave sleep (SWS). This refers to the phenomenon that during sleep, hippocampal cells activate a specific pattern during sleep, which seems to be similar to the activation during encoding (Wilson & McNaughton, 1994). Although causal evidence is currently missing, this reactivation correlates with memory performance (Derdikman & Moser, 2010; Dupret, O’neil, Pleydell-Bouverie, & Csicsvari, 2010). Therefore, it seems to reflect memory consolidation. This reactivation can be induced by using sensory cues (Carr, Jadhav, & Frank, 2011). Thus, TMR seems to make use of the hippocampal replay by activating sensory cues associated with previous learning and therefore enhances learning.

A typical TMR paradigm can be divided into four phases according to Cellini & Capuozzo (2018). The first phase is the encoding phase, in which participants learn new material. This new material is learned while it’s being associated with sensory cues (typically auditory or olfactory). After the encoding phase, participants are given a memory test to see how well they remembered the learned material. Then, participants are instructed to go to sleep. While sleeping the participants are re-exposed to half of the associated sensory cues. After sleeping, the participants are tested again on their memory of the learned material. What is often found is that cued items are better remembered than uncued items.

The type of modalities used to associate the learned material with are typically olfactory or auditory. The use of olfactory stimuli has several benefits for TMR paradigms. Odors are strong at eliciting contextual retrieval cues for declarative and emotional memory (Schouten et al., 2017). Besides that odors rarely awaken sleepers, making them thus very useful for TMR paradigms (Rasch, Büchel, Gais, & Born, 2007). Auditory stimuli on the other hand are more arousing than odors (Oudiette & Paller, 2013; Schouten et al., 2017) However, they have been more successful than olfactory cues in various domains, including procedural memories. Schouten et al (2017) hypothesize that olfactory stimuli are best suited for declarative and emotional memories, while auditory can be best used for procedural memories. Auditory cues have the advantage of the ability to be congruent with the learning material (for example, in a TMR paradigm that uses a word learning task, the words can be re-exposed in sleep by verbally presenting the participants with the cued words). Congruent learning has been found to be more effective than incongruent learning (Oudiette & Paller, 2013; Cairney, Lindsay, Sobczak, Paller, & Gaskell, 2016; Groch et al., 2016).

Besides modality, the effect of TMR is dependent on several variables. Firstly, TMR cannot enhance memories when the material is already perfectly learned. Inversely, when participants haven’t properly encoded the learning material, they won’t benefit from subsequent cueing (Schouten et al., 2017). Additionally, State of consciousness matters, as it seems that TMR during sleep has been more successful than TMR during wake. This is illustrated by Diekelmann, Büchel and Rasch (2011), who used a TMR paradigm with olfactory cues and an object-location task. They found that participants that received TMR during wake showed decreased performance on the post-nap object-location task, while the sleep group showed the opposite results. They hypothesize that cueing while awake destabilizes the memory, and makes it prone to interference, while cueing in sleep immediately stabilizes memories and thus makes them resistant to interference. Further, time spend in SWS seems to be predictive of how strong the TMR effect is (Cariney, Durrant, Hulleman, & Lewis, 2014). Age also seems to influence the effect of TMR. Cordi, Schreiner and Rasch (2018) used a TMR paradigm with a Dutch vocabulary learning task with older adults as participants. They found that TMR enhanced performance for younger adults, yet not for older adults. They reason that this could be due to decrease in hippocampal replay mechanism during SWS. It also matters in what way you present the cue during initial learning. Schouten et al., (2017) Mentions that both auditory and olfactory cues have the chance of being too ordinary which prevents them from become meaningful in associating with the learned material. Additionally, both types are prone to sensory habituation and should thus not be presented too often.

Most studies of TMR focus on finding out under what circumstances TMR is beneficial. Besides that, there are also several reviews that focused on evaluating the literature of TMR. Cellini & Capuozzo (2018) reviewed TMR studies on whether the results depend on type of memory, task employed, sensory cue used and sleep stage of stimulation. Their review is a comprehensive summary of the literature, yet it lacks systematic methods which leads to a chance of not incorporating all studies on TMR. Schouten et al (2016) provide a similar overview of the literature, focusing on giving an overview of the limitations and pitfalls of using TMR. Lastly, Oudiette and Paller (2013) focused their review on describing the method of TMR and attempt to provide new perspectives on how sleeping contributes to memory. All these reviews succeed at what they aimed to do, yet none have focused yet on systematically evaluating the size of the effect of TMR.

The aim of this study was to conduct a meta-analysis on the effect of TMR. Besides the main aim, this study also aimed at evaluating whether the effect of TMR is dependent on modality, sleep stage and type of memory. Modality most likely depends on whether a study that used auditory TMR uses congruent signals. However, most studies using auditory TMR don’t use congruent signals, therefore it is hypothesized that TMR is not influenced by modality. Furthermore, TMR has been successfully tested for procedural, semantic and spatial memory. Yet emotional memory shows mixed results (Schouten et al., 2017; Cellini & Capuozzo, 2018). Therefore, it is hypothesized that the effect of TMR depends on whether the memory tested is emotional or not. Study quality will also be taken into account to evaluate the quality of the evidence. To attain these aims, a comprehensive literature search was carried out. The extracted data was then analyzed using a random effects model, as this is more suitable than a fixed model for analyzing studies that employed different methodology. Study heterogeneity was assessed. Publication bias was assessed by inspecting a funnel plot. Furthermore, subgroup analyses were carried out on modality and type of memory. Sleep stage wasn’t possible as most studies induced TMR during SWS.

## Method

### Identification of studies

A comprehensive literature search was carried in Pubmed, Psychinfo and Web of Science, to find experimental studies that tested the effect of targeted memory reactivation on recall. The time interval used for all databases was the same (from 01-01-2000 till 20-02-2019). The search query used was “targeted memory reactivation OR (memory reactivation AND sleep) OR (cue* AND memory AND sleep) NOT cued recall”, and the search focused on title, abstract and keywords. To supplement the literature search in the databases, several articles found by examining reference lists of articles about targeted memory reactivation.

### Inclusion exclusion criteria

Study eligibility was determined on several inclusion and exclusion criteria. Inclusion criteria were: (1) experimental studies only, (2) available in English, (3) containing healthy human participants, (4) targeted memory reactivation must occur during sleep. The exclusion criteria were: (1) no full text available and (2) no dissertations.

### Data collection and assessment

Data was collected on sample size, age range (or if range was not reported, the mean age), the method, sleep stage that TMR was introduced in, modality exposed to TMR, type of memory, significance (in either positive or negative direction). It was decided to categorize type of memory quite broadly, as often studies did not specify the type of memory they tested or used interchangeable terms. Further, and most relevant for this meta-analysis, the mean, standard error of the mean or standard deviation and sample size used in the analysis were collected for the experimental and control group.

Furthermore, The methodological quality of each included study was assessed using the Downs and black (1998) checklist, which can be used to assess randomized studies. The checklist consists 27 questions which are divided in 5 subscales. All questions are scored either 1 or 0 (with the exception of 1 question, which can be also scored 2), thus the maximum score that can be attained is 28. The subscales are respectively: (1) reporting (n=10), (2) external validity (n=3), (4) internal validity - bias (n=6), (5) internal validity - confounding (n=6) and (7) power (n=1). The quality scores were divided in good (scores > 21), moderate (scores ranging 11 - 20) and low (scores < 10) (Hartling, Brison, Crumley, Klassen, & Pickett, 2004).

### Analysis

Hedge’s g was computed using the extracted data. A random effects model was chosen to estimate the pooled effect sizes. The DerSimonian-Laird method was used to estimate tau^2^. Most TMR studies differ on method, modality and memory system tested. Therefore, it was decided to use an random effects model which assumes that besides variation due to sampling error, another source of variance in the data. This other source of error in this case was the difference in method, modality and memory system tested. Furthermore, two subgroup analyses were performed to check whether the Hedges g differed based on modality and memory. Initially it was also planned to conduct a subgroup analysis on sleep stage, however the TMR studies almost exclusively introduced TMR during slow wave sleep. Therefore it was decided to leave out the only REM study and not conduct a subgroup analysis on this variable. To further explore the data, an influence analysis, a Baujat plot and leave-one-out-analyses were computed to see whether certain studies had a large influence on the results of the random effects model. All analyses were conducted using the dplyr, meta and metafor libraries in Rstudio (Wickham, Francois, Henry, & Müller, 2015; Schwarzer & Carpenter, 2010; Viechtbauer, 2010).

## Results

### Literature search and excluded studies

The initial search results were 264 studies for Pubmed, 374 studies for Psychinfo and 229 for Web of Science. From these search results 121 were initially identified as relevant to TMR. On more careful inspection of the articles, 76 were excluded for either using TMR, being review articles or dissertations, making use of TMR. One study was excluded for supplementing the TMR procedure used with drugs. Thus in total, 41 studies were deemed relevant for further examination.

### Methodological quality

The 41 studies deemed relevant were assessed on their methodological quality by one judge (this limitation is further explained in the discussion). The average study quality was (m=21.7, SD = 2.1) and was thus considered as good. The quality scores ranged from 17 to 26. About 70.7% of the studies were considered good, the remaining 29.3% were considered moderate. The means per subscale were respectively: Reporting 9 out of 11, external validity 1.4 out of 3, internal validity 5.5 out of 7, and selection bias 5.2 out of 6.

### Study characteristics

In 29 of the 41 studies (69%), TMR was induced during slow wave sleep. The other 12 studies did either: not specify the sleep stage (n=4), used NREM2 and SWS (n=4), used NREM2 (n=2), used REM and NREM2 (n=1) or used REM (n=1). Due to the lack of variation in sleep stage, it was decided to not conduct a subgroup analyses for this variable. For modality, 31 out of 41 were auditory stimuli (75.6%), the remaining 10 studies were olfactory. Memory was respectively categorized in to the following categories: Emotional (n=4), procedural (n=8), semantic (n=15) and spatial (n=14). Out of the 41 studies, 26 were significant (63.4%). Out of all the studies, 25 reported enough descriptive statistics to be used for the random effects model. The proportion of significance between the studies that did report descriptive (61.5%) did not seem different than studies that did not (66.67%). out of the 26 studies used in the analyses, 7 were olfactory and 18 were auditory. Further, 3 were emotional, 4 were procedural, 10 were semantic and the remaining 8 were spatial. The full overview is presented in Appendix A.

### Effect of TMR

This random effects model yielded an overall controlled effect size of hedges g = 0.4150 (95% CI= 0.2691 - 0.5609, t = 5.87, p < 0.0001), which suggests that TMR has a moderate to small effect on memory enhancement. The prediction interval ranged from - 0.1461 - 0.9752, which indicates that the effect of TMR is not robust in every context. The measures of heterogeneity yielded the following results: tau^2^=0.0686, I^2^=10.3% (95% CI= 0% - 43.6%), Q = 26.76, df = 24, p=0.3158. indicating that a relatively high amount of variance is due to sampling error, not true effect variance. Taken together, the measures of heterogeneity do not seem to indicate that the variance of the studies is largely due to variance in true effects.

### Influence analyses

To see whether the fit of the model could be increased, several influence analyses were conducted to check whether some studies had a large influence on between-study heterogeneity. Firstly, inspecting the influence analyses measures, none of the studies indicate significant bias (Appendix C, studies in red indicate possible influential studies). The baujat plot, indicates that Schreiner, Lehmann, and Rasch (2015) could be an influential study (Appendix D). Furthermore, the leave-one-out-analyses in which the I^2^, confidence interval of the hedges g and estimated effect is recalculated with one study omitted each time indicates that leaving out one of the studies does not decrease the I^2^ by a substantial amount Appendix E). Thus none of the studies seem to influence the model substantially

### Subgroup analysis: Modality

The test for subgroup differences between modalities was non-significant (Q=0.29, df=1, p=0.5888). This indicates that the effect of TMR was not dependent on the modality used. The heterogeneity measures for auditory were Q=18.21, tau^2^=0.0582 and I^2^=.6.6%. For olfactory they were Q=8.50, tau^2^=0.1013 and I^2^=29.4%. Confidence interval was auditory (95% CI = 0.2296 - 0.5586) and olfactory (95% CI = 0.0859 - 0.9012). So even though the model yielded a non-significant result, the CI for auditory seems more precise, which might indicate that the effect of TMR is more robust for auditory TMR. Forest plot is presented in appendix G.

### Subgroup analysis: Memory

The test for subgroup differences between memory systems was significant (Q=35.62, df=3, p<0.0001). This indicates that the effect of TMR depends on what kind of memory you reactivate. The heterogeneity measures were emotional (Q=0.03, tau^2^<0.0001, I^2^=0%) procedural (Q=1.64, tau^2^=0.0187, I^2^=0%), semantic (Q=13.17, tau^2^=0.0834, I^2^=31.7%) and spatial (Q=6.30, tau^2^=0.0633, I^2^=0%). The confidence intervals were Emotional (95% CI = - 0.0370 - 0.1613), procedural (95% CI= 0.2180 - 1.0291), semantic (95% CI= 0.1189 - 0.6606) and spatial (95% CI=0.2030 - 0.7904). The effects of emotional were the lowest (g=0.0621) while the effects of procedural were the highest (g=0.6235). However these results should be cautiously interpreted due to the low sizes of the subgroups (emotional n = 3, procedural n = 4, semantic n = 10, spatial n = 8) and therefore most likely lack power to detect meaningful differences. Forest plot is presented in appendix H.

### Publication bias

A funnel plot was inspected to determine if there was potential publication bias. Upon inspecting the funnel plot, it seemed that the plot is not fully symmetrical, as well that the studies aren’t lined up in the shape of the funnel. The reason why the studies aren’t shaped like the funnel could be that the variance of the sample sizes was quite low, which could explain why the standard error thus did not differ a lot between studies. The funnel seemed to be asymmetrical in the bottom-left corner, indicating that small studies with low effect sizes are missing. Therefore, the results of the random effects models should be interpreted cautiously, as they could be influenced by publication bias. The funnel plot is presented in appendix I.

## Discussion

The aim of this study was to conduct a meta-analysis on the effect of TMR. The Random effects model yielded a significant effect, with a small to moderate effect size. This indicates that the effect of TMR is successful at enhancing memories while sleeping. Furthermore the subgroup analyses yielded a significant result for memory, yet not for modality. The effect of TMR is thus dependent on what type of memory tested. Emotional memories benefited the least from TMR, while spatial memories seemed to benefit the most. The subgroup analyses of memory however should be interpreted with caution, as the amount of studies per sub group were quite low, and thus might lack power to detect meaningful differences.

As hypothesized, modalities did not significantly differ on their effect size of TMR. Currently no study has directly compared auditory to olfactory cues, making it hard to give any informed reasons on why this is the case. Several studies do indicate that memories congruent with the sensory cues were easier learned than others (Cairney et al, 2016; Groch et al., 2016). According to these results, it therefore seems that when cues are incongruent with the learned material, one is preferred over the other. However, as congruent cues can elicit a stronger TMR effect, it makes sense that auditory cues congruent with learning material are strongest. Thus, there could be an interaction effect between modality and memory. This could not be tested in this study due to the lack of studies available and thus lack of power.

Type of memory was hypothesized to be dependent on whether the memories tested were emotional or not. Looking at the forest plot for memory (appendix I) it seems to be that emotional memories are not influenced by TMR, while other effects generally are similar to each other. Schouten et al., (2017) as well Cellini and Capouzzo (2018) describe that the TMR effect on emotional memory shows mixed results. Both describe that it is unclear whether NREM or REM sleep is best for cueing emotional memories. Furthermore, previous behavioural studies show that emotional memory consolidation is generally stronger than neutral memory consolidation during sleep, especially during REM sleep (Cunningham, Pardilla-Delgado, Alger, & Payne, 2014). However, one study that used TMR during REM sleep did not find that TMR had a different effect for neutral and emotional stimuli (Rihm & Rasch, 2015). This might mean that emotional memories might be more resistant to external interference, thus TMR might not be beneficial for enhancing these types of memories. Currently however, it is difficult to make hard conclusions based on the little data there was on emotional memories.

The results indicated that there was no substantial heterogeneity in the data. However, the effect has a large prediction interval, indicating that the effect of TMR could be dependent on memory or task. Task wasn’t tested here, if it was it would have most likely lacked power due to the high variety of tasks employed in TMR studies. Furthermore, similar tasks could also differ on what they actually measure therefore it is hard to study how task influences TMR. Memory types vary less, yet there were little studies per subgroup, especially emotional and procedural memory hasn’t been tested a lot. The funnel plot indicates that some publication bias might be presented, and this would mean that the effect presented here is overestimated. Sterne et al., (2011) describe several reasons why funnel plots are asymmetrical. Alternative reasons for asymmetry can be ascribed to poor methodological quality (poor design, inadequate analysis, or fraud), True heterogeneity (size of the effects differs according to study size), and by chance. Firstly chance seems unlikely due to the number of studies incorporated. Poor methodological quality also seems unlikely as the average quality was considered good. Heterogeneity was quite low, so this most likely wouldn’t have influenced the funnel plot. Therefore, the lack of symmetry in the bottom half of the funnel plot might be due to publication bias.

The study characteristics indicated that most studies focused on applying TMR during SWS. This was mostly due to a lack of precision in employing TMR (TMR is applied during NREM 2 and SWS because some participants did not reach SWS), not defining sleep stage. That means that relatively little studies have tried to find out what the benefits of cueing in REM and NREM 2 are. Therefore it was decided to leave REM sleep out of the analysis. The two studies included in this study who cued specifically during REM sleep both did not yield significant effects (Rihm et al., 2015; Cordi et al., 2014). Besides that, it is as of now uncertain whether TMR during REM sleep has any effect, even though REM sleep does play a role in memory consolidation (Cunningham et al., 2014). Cordi et al., (2014) theorize that their lack of effect could be because even though memory reactivations exist during REM sleep, they don’t have a functional effect on stabilizing memory consolidation. Rihm et al., (2015) assumed that REM sleep might be prone to cueing emotional memories. They found that cueing influenced reduced arousal, yet did not affect memory strength. The conclusion of Cordi et al. is in line with these findings. REM sleep might have an important role on memory, but not necessarily on memory consolidation

Although the study quality was on average high, the external validity was low. Often, the study incorporated students, who were highly educated and young. If studies did not report whether they sampled students, they often did not report education level or other principal confounders. Hanel and Vione (2016) found that generalizing student samples to the general public can be problematic when they differ on personal variables, like intelligence. In the context of TMR, it is hard to generalize the results further than students as intelligence is related to how well memories are consolidated during sleep (Bódizs et al., 2005; Schabus et al., 2006). Further, students are often young adults or adolescents. One study using found TMR might not be beneficial for older adults, indicating that TMR might be age dependent (Cordi et al 2018). The results of this study should therefore be carefully interpreted when generalizing to populations other than students. Further studies should focus on testing whether intelligence matters for the effect of TMR as well as testing how the effect of TMR is dependent on age.

This meta-analysis suffers from several limitations. Firstly, not every study was taken into the analysis due to a lack of access to descriptive statistics needed to conduct the analysis. This influences the precision of the effect. The size of the effect could be underestimated, as the proportion of significant results was slightly higher (66.67%) for the non-included studies than the included studies (61.54%). Another limitation is that the subgroup analysis for memory lacks power due to the fact that there weren’t a lot of studies testing certain type of memories (procedural and emotional). Further studies could thus be done for these types of memories as research is lacking in that area. Lastly, The study quality was scored by only one individual, while usually this is done by at least two to calculate an inter-reliability coefficient. Without comparison to another judge, it is hard to determine whether there was experiment bias, therefore the results of the methodological quality should be interpreted with caution.

To conclude, this study is the first to evaluate the effect of TMR. It provides evidence that TMR has the ability to enhance memories is weak to moderate. This effect is independent of modality but does depend on the type of memory. There are no indications of substantial study heterogeneity. Results might have been influenced by publication bias. Further studies should find out whether TMR is dependent on age and intelligence, as both these factors could determine how generalizable the results are. Furthermore, even though TMR has the potential to be of practical use, there is a lack of studies that try to induce TMR outside of the laboratory. Since the data collection for this study, one study has been published researching this, which successfully showed that TMR unsupervised at home can enhance memory (Göldi & Rasch, 2019). More studies however are needed to confirm these results.

## Appendix A: Study Characteristics and Study Quality

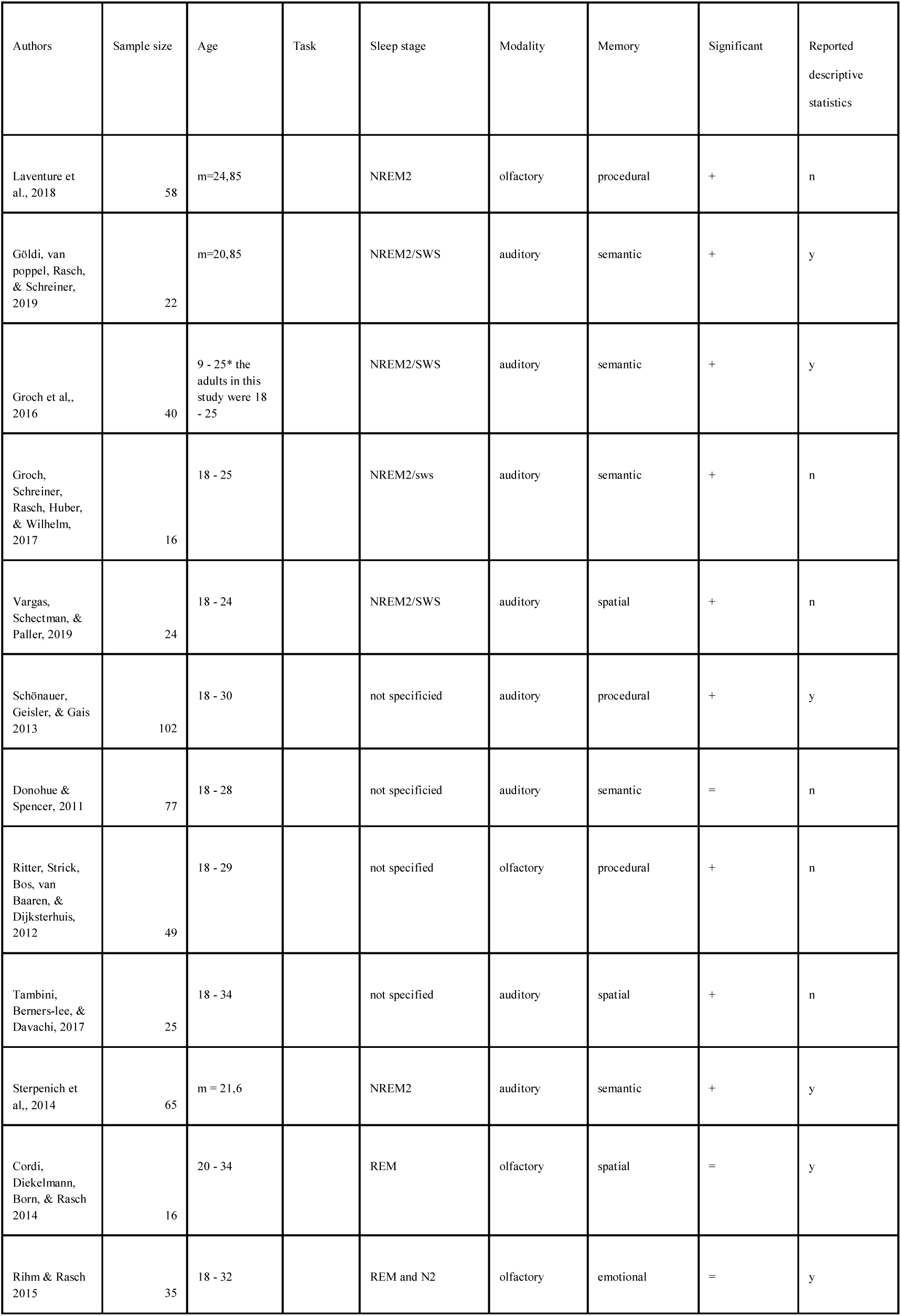
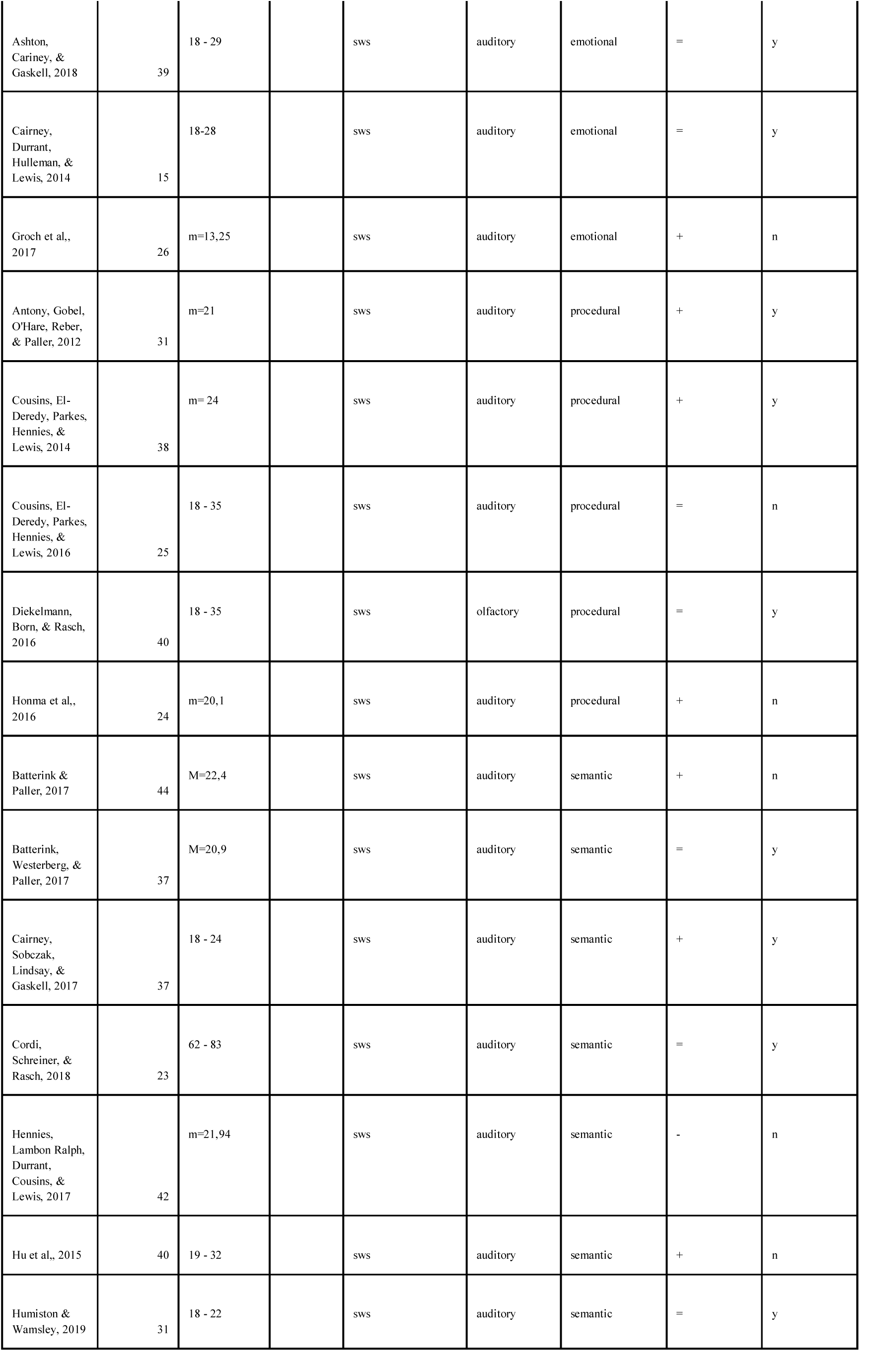
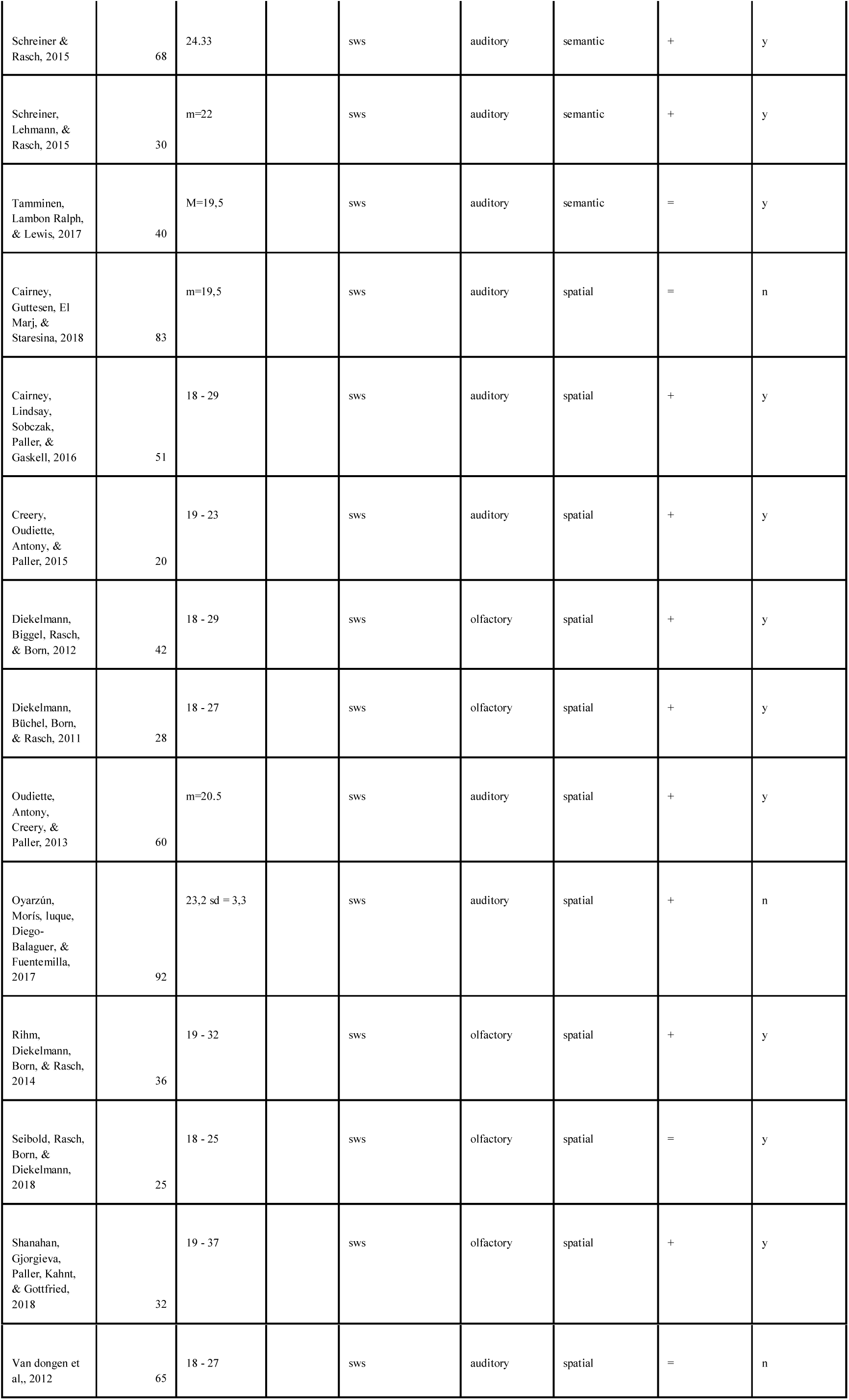

| Authors | Sample size | Age | Task | Sleep stage | Modality | Memory | Significant | Reported descriptive statistics |
| --- | --- | --- | --- | --- | --- | --- | --- | --- |
| Laventure et al., 2018 | 58 | m=24,85 |  | NREM2 | olfactory | procedural | + | n |
| Göldi, van poppel, Rasch, & Schreiner, 2019 | 22 | m=20,85 |  | NREM2/SWS | auditory | semantic | + | y |
| Groch et al., 2016 | 40 | 9 - 25* the adults in this study were 18 - 25 |  | NREM2/SWS | auditory | semantic | + | y |
| Groch, Schreiner, Rasch, Huber, & Wilhelm, 2017 | 16 | 18 - 25 |  | NREM2/sws | auditory | semantic | + | n |
| Vargas, Scheetman, & Paller, 2019 | 24 | 18 - 24 |  | NREM2/SWS | auditory | spatial | + | n |
| Schönauer, Geisler, & Gais 2013 | 102 | 18 - 30 |  | not specified | auditory | procedural | + | y |
| Donohue & Spencer, 2011 | 77 | 18 - 28 |  | not specified | auditory | semantic | = | n |
| Ritter, Strick, Bos, van Baaren, & Dijksterhuis, 2012 | 49 | 18 - 29 |  | not specified | olfactory | procedural | + | n |
| Tambini, Berners-lee, & Davachi, 2017 | 25 | 18 - 34 |  | not specified | auditory | spatial | + | n |
| Sterpenich et al., 2014 | 65 | m = 21,6 |  | NREM2 | auditory | semantic | + | y |
| Cordi, Diekelmann, Born, & Rasch 2014 | 16 | 20 - 34 |  | REM | olfactory | spatial | = | y |
| Rihm & Rasch 2015 | 35 | 18 - 32 |  | REM and N2 | olfactory | emotional | = | y |
| Ashton,<br>Carney, &<br>Gaskell, 2018 | 39 | 18 - 29 |  | sws | auditory | emotional | = | y |
| Cairney,<br>Durrant,<br>Hulleman, &<br>Lewis, 2014 | 15 | 18-28 |  | sws | auditory | emotional | = | y |
| Groch et al.,<br>2017 | 26 | m=13,25 |  | sws | auditory | emotional | + | n |
| Antony, Gobel,<br>O'Hare, Reber,<br>& Paller, 2012 | 31 | m=21 |  | sws | auditory | procedural | + | y |
| Cousins, El-Deredy, Parkes,<br>Hennies, &<br>Lewis, 2014 | 38 | m= 24 |  | sws | auditory | procedural | + | y |
| Cousins, El-Deredy, Parkes,<br>Hennies, &<br>Lewis, 2016 | 25 | 18 - 35 |  | sws | auditory | procedural | = | n |
| Diekelmann,<br>Born, & Rasch,<br>2016 | 40 | 18 - 35 |  | sws | olfactory | procedural | = | y |
| Honma et al.,<br>2016 | 24 | m=20,1 |  | sws | auditory | procedural | + | n |
| Batterink &<br>Paller, 2017 | 44 | M=22,4 |  | sws | auditory | semantic | + | n |
| Batterink,<br>Westerberg, &<br>Paller, 2017 | 37 | M=20,9 |  | sws | auditory | semantic | = | y |
| Cairney,<br>Sobczak,<br>Lindsay, &<br>Gaskell, 2017 | 37 | 18 - 24 |  | sws | auditory | semantic | + | y |
| Cordi,<br>Schreiner, &<br>Rasch, 2018 | 23 | 62 - 83 |  | sws | auditory | semantic | = | y |
| Hennies,<br>Lambon Ralph,<br>Durrant,<br>Cousins, &<br>Lewis, 2017 | 42 | m=21,94 |  | sws | auditory | semantic | - | n |
| Hu et al., 2015 | 40 | 19 - 32 |  | sws | auditory | semantic | + | n |
| Humiston &<br>Wamsley, 2019 | 31 | 18 - 22 |  | sws | auditory | semantic | = | y |
| Schreiner & Rasch, 2015 | 68 | 24.33 |  | sws | auditory | semantic | + | y |
| Schreiner, Lehmann, & Rasch, 2015 | 30 | m=22 |  | sws | auditory | semantic | + | y |
| Tamminen, Lambon Ralph, & Lewis, 2017 | 40 | M=19,5 |  | sws | auditory | semantic | = | y |
| Cairney, Guttesen, El Marj, & Staresina, 2018 | 83 | m=19,5 |  | sws | auditory | spatial | = | n |
| Cairney, Lindsay, Sobczak, Paller, & Gaskell, 2016 | 51 | 18 - 29 |  | sws | auditory | spatial | + | y |
| Creery, Oudiette, Antony, & Paller, 2015 | 20 | 19 - 23 |  | sws | auditory | spatial | + | y |
| Diekelmann, Biggel, Rasch, & Born, 2012 | 42 | 18 - 29 |  | sws | olfactory | spatial | + | y |
| Diekelmann, Büchel, Born, & Rasch, 2011 | 28 | 18 - 27 |  | sws | olfactory | spatial | + | y |
| Oudiette, Antony, Creery, & Paller, 2013 | 60 | m=20.5 |  | sws | auditory | spatial | + | y |
| Oyarzún, Morís, luque, Diego-Balaguer, & Fuentemilla, 2017 | 92 | 23,2 sd = 3,3 |  | sws | auditory | spatial | + | n |
| Rihm, Diekelmann, Born, & Rasch, 2014 | 36 | 19 - 32 |  | sws | olfactory | spatial | + | y |
| Seibold, Rasch, Born, & Diekelmann, 2018 | 25 | 18 - 25 |  | sws | olfactory | spatial | = | y |
| Shanahan, Gjorgieva, Paller, Kahnt, & Gottfried, 2018 | 32 | 19 - 37 |  | sws | olfactory | spatial | + | y |
| Van dongen et al., 2012 | 65 | 18 - 27 |  | sws | auditory | spatial | = | n |

## Appendix B: Forest Plot Random Effects Model

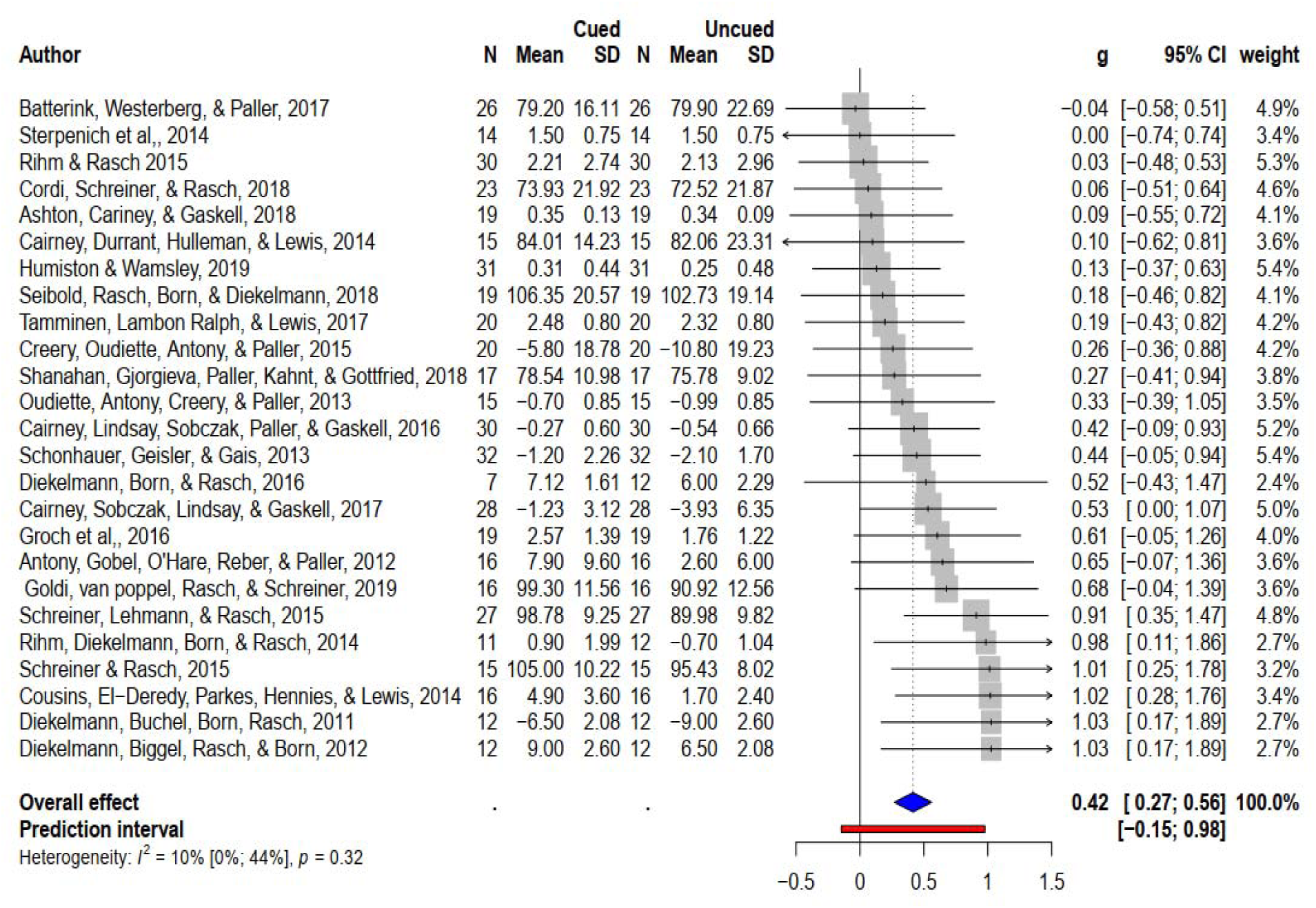

## Appendix C: Influence Analyses

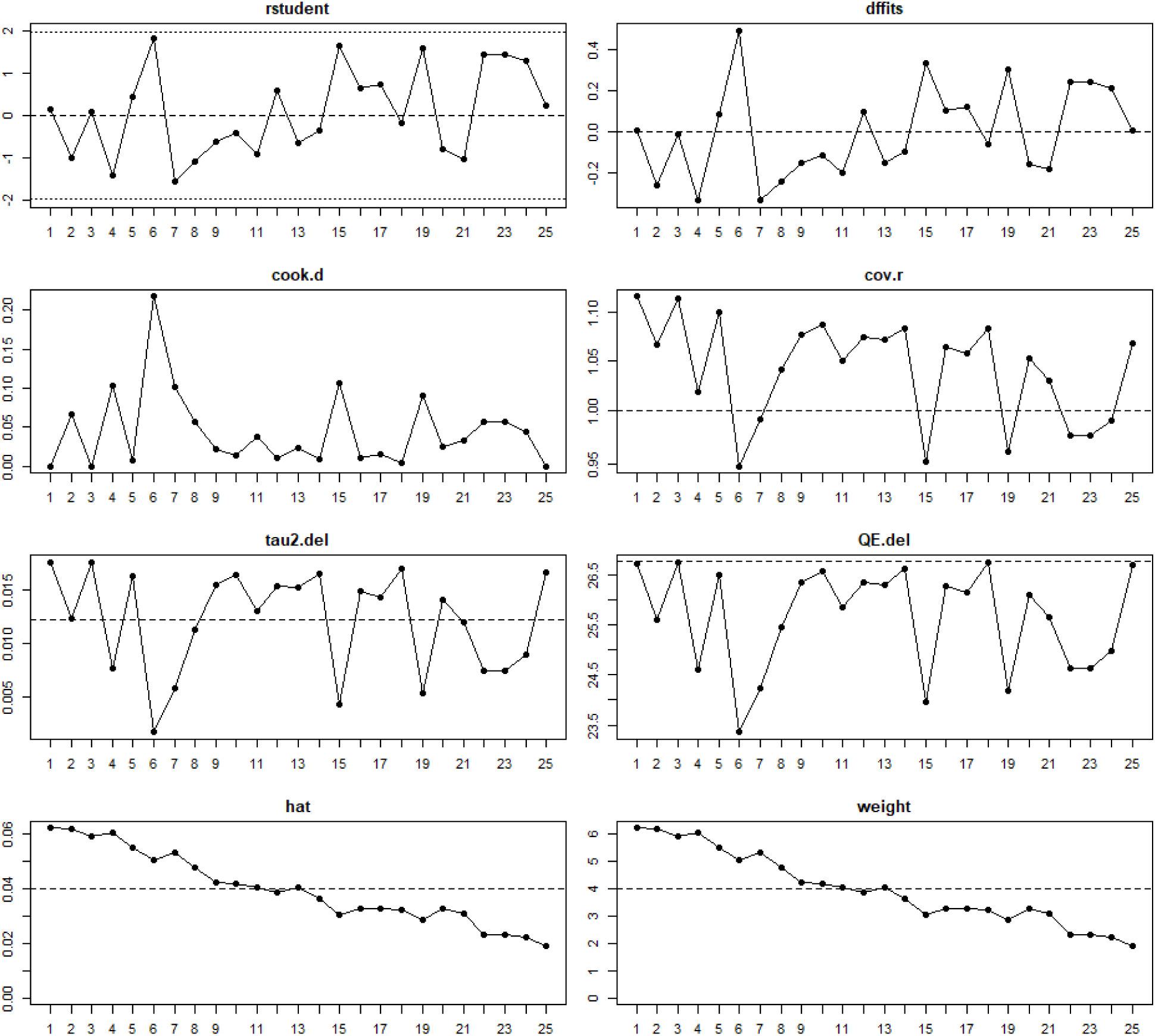

## Appendix D: Baujat plot

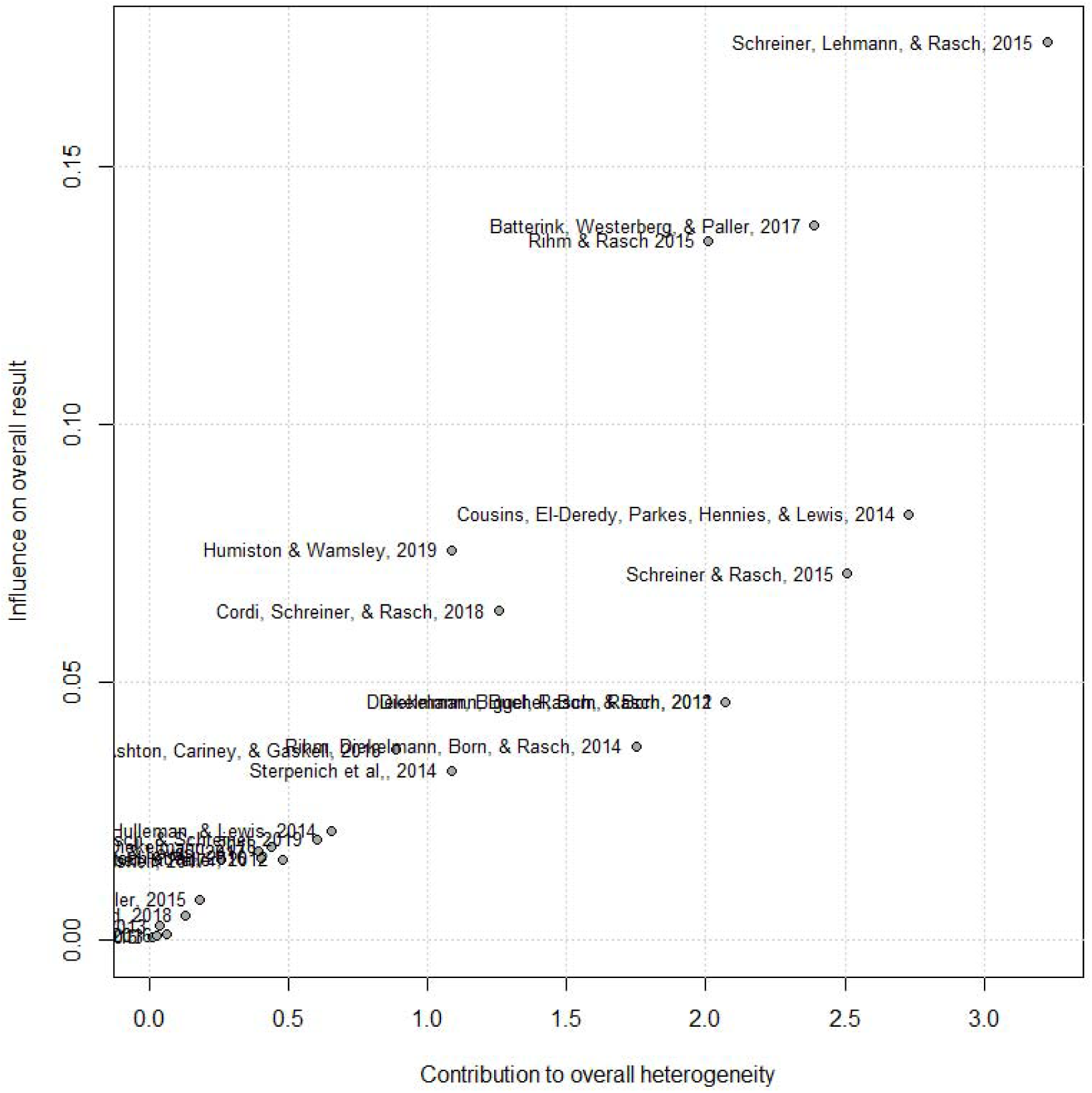

## Appendix E: Leave-one-out analysis

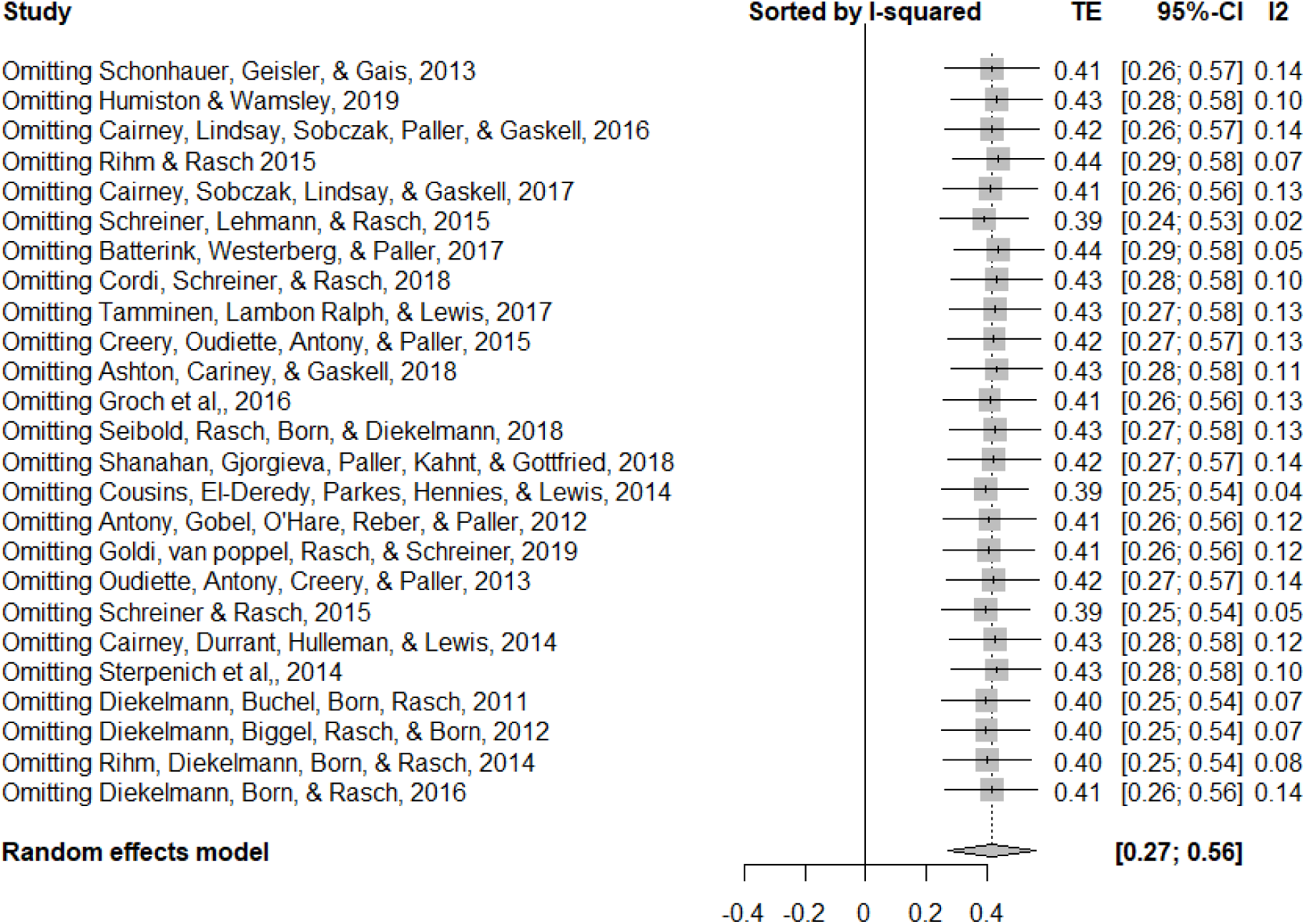

## Appendix F: Subgroup analysis plot – Modality

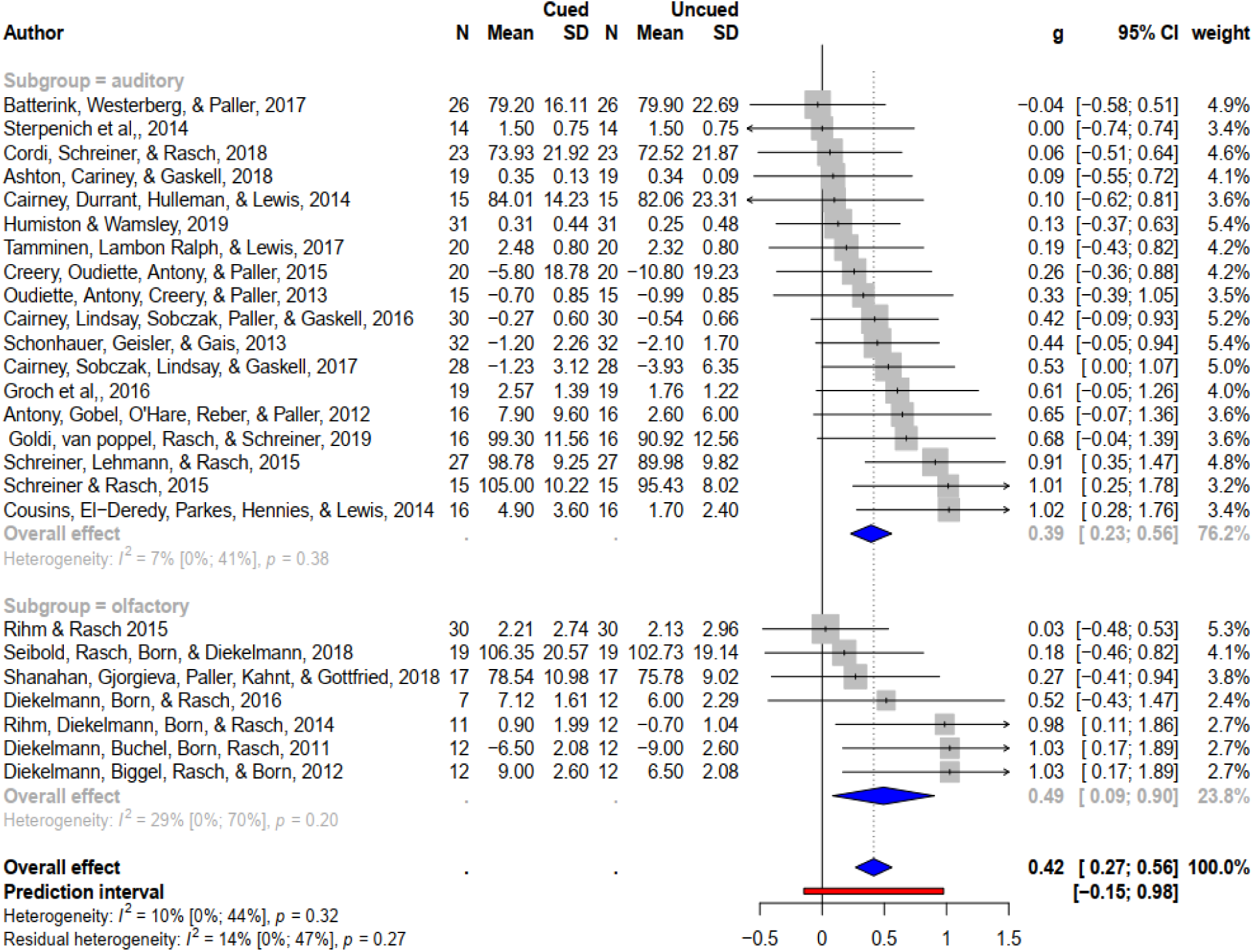

## Appendix G: Subgroup analysis plot – Memory

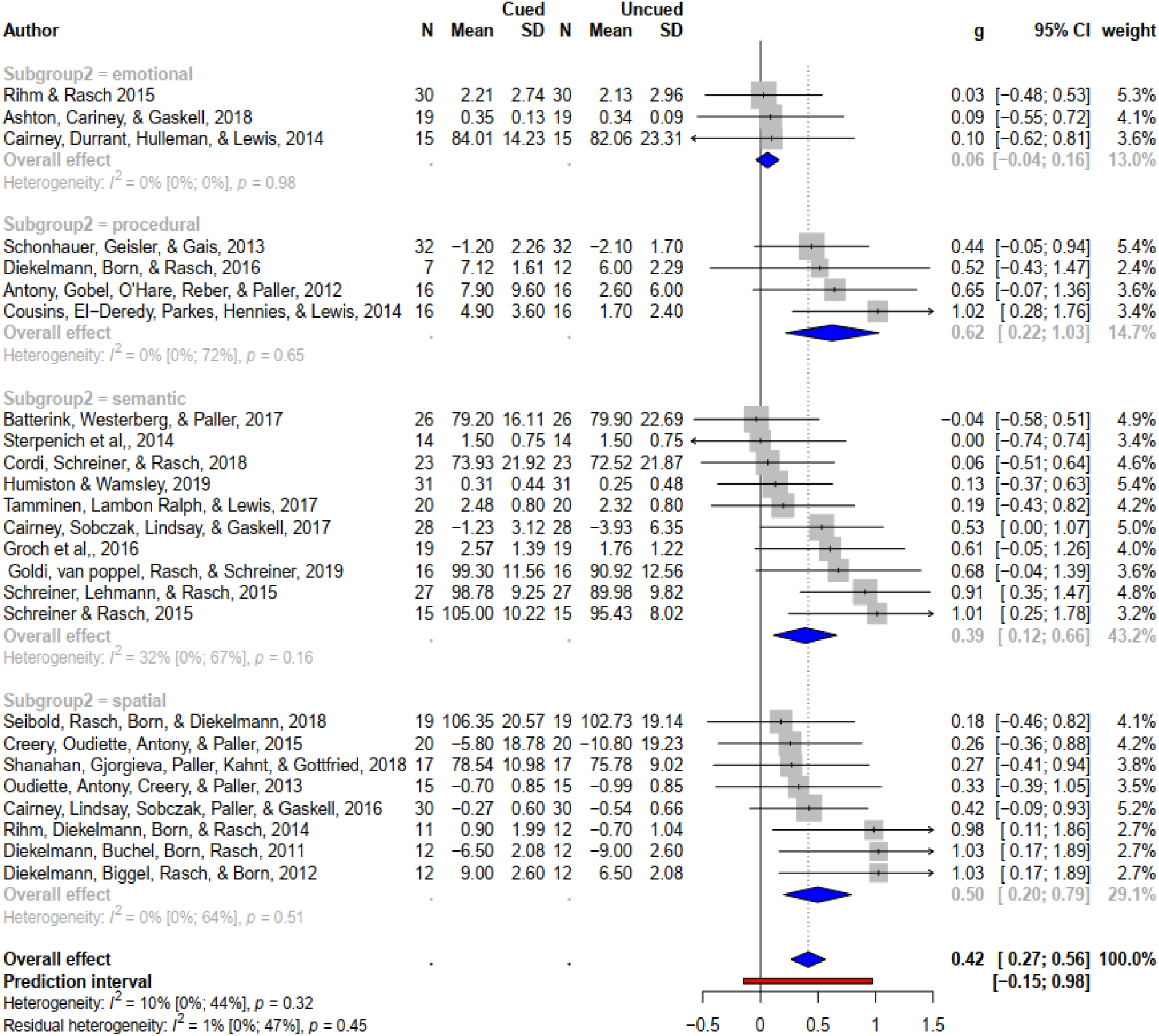

## Appendix H: Funnel plot

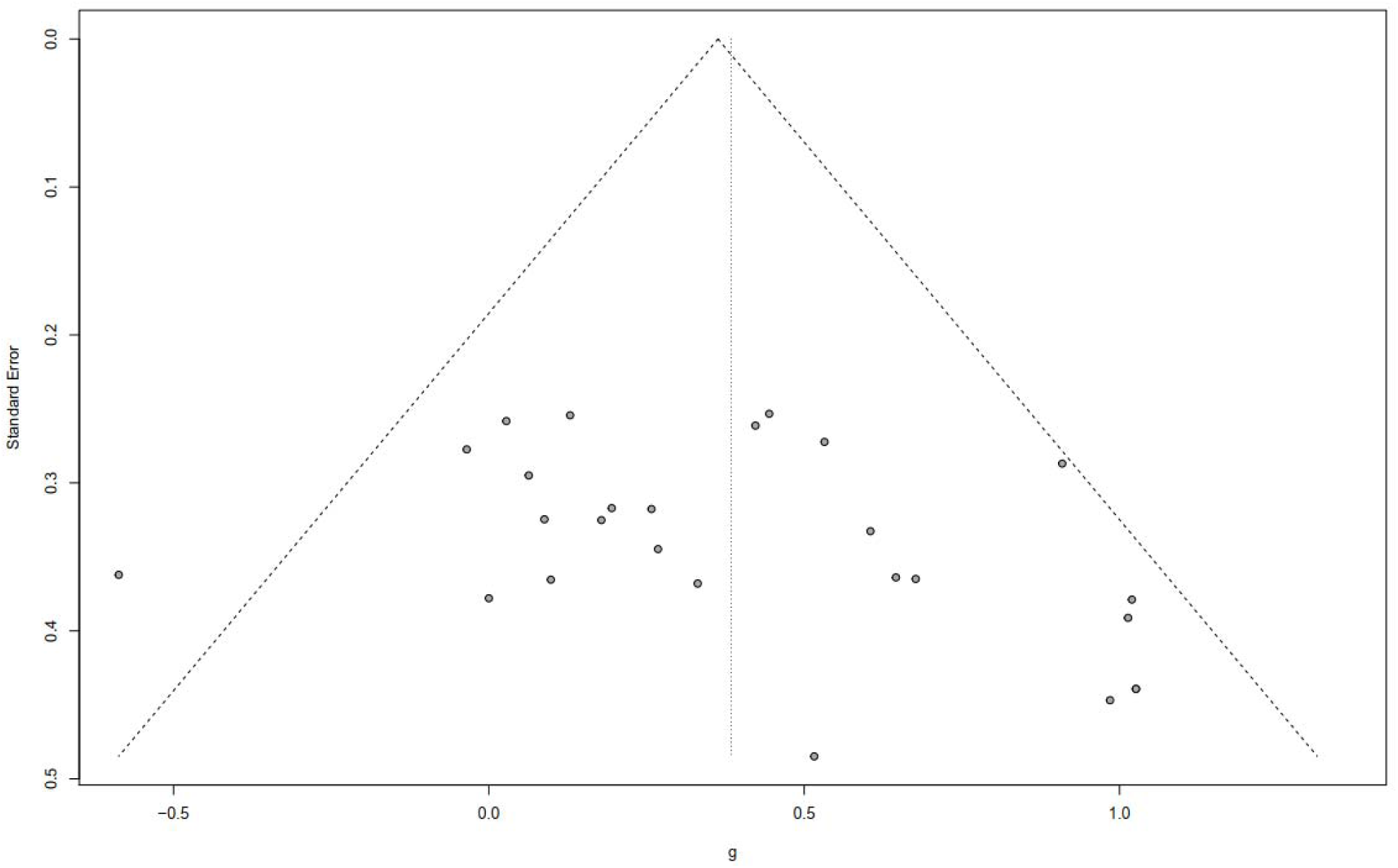

